# Selective Inhibition of Cytosolic PARylation via PARG99: A Targeted Approach for Mitigating FUS-associated Neurodegeneration

**DOI:** 10.1101/2024.11.25.625276

**Authors:** Manohar Kodavati, Vikas H. Maloji Rao, Joy Mitra, Muralidhar L. Hegde

**Author notes:** Corresponding authors: M.K.; M.L.H. Corresponding authors’ address: Manohar Kodavati, PhD Research Associate, Division of DNA Repair Research Center for Neuroregeneration, Department of Neurosurgery Houston Methodist Research Institute (HMRI) Houston, Texas 77030, USA. Muralidhar L. Hegde, PhD Professor, Department of Neurosurgery Everett E. and Randee K. Bernal Centennial Endowed Chair of Neurological Institute Director, Division of DNA Repair Research, Center for Neuroregeneration Houston Methodist Research Institute 6670 Bertner Ave, RI-11-112.

## Abstract

Neurodegenerative diseases such as amyotrophic lateral sclerosis (ALS) are characterized by complex etiologies, often involving disruptions in functions of RNA/DNA binding proteins (RDBPs) such as FUS and TDP-43. The cytosolic mislocalization and aggregation of these proteins are linked to accumulation of unresolved stress granules (SGs), which exacerbate the disease progression. Poly-ADP-ribose polymerase (PARP)-mediated PARylation plays a critical role in this pathological cascade, making it a potential target for intervention. However, conventional PARP inhibitors are limited by their detrimental effects on DNA repair pathways, which are already compromised in ALS. To address this limitation, we investigated a strategy focused on targeting the cytosolic compartment by expressing the cytosol-specific, natural PAR-glycohydrolase (PARG) isoform, PARG99. Using ALS patient derived FUS mutant induced pluripotent cells (iPSCs) and differentiated neurons, we observed elevated levels of FUS in insoluble fractions in mutant cells compared to mutation-corrected isogenic lines. The insoluble FUS as well as TDP-43 levels increased further in sodium arsenite-treated or oxidatively stressed cells, correlating with accumulation of unresolved SGs. Notably, both PARG99 and PARP inhibitors reduced SG formation and insoluble FUS levels, however, PARG99 treated cells exhibited significantly lower DNA damage markers and improved viability under oxidative and arsenite stress. This study highlights the potential of PARG99 as a cytosol-specific intervention to mitigate FUS-associated toxicity while preserving critical nuclear DNA repair mechanisms, offering a promising strategy for addressing the underlying pathology of ALS and potentially other SG-associated neurodegenerative diseases.

## Introduction

ADP-ribosylation is a conserved, reversible post-translational modification (PTM) involving the addition of ADP-ribose moieties to target proteins, using nicotinamide adenine dinucleotide (NAD+) as a substrate [1]. This modification plays a crucial role in various cellular processes, including DNA repair, stress response, signal transduction, and apoptosis [2]. ADP-ribosylation can occur as either mono ADP-ribosylation (MARylation) or poly ADP-ribosylation (PARylation), with the latter being catalyzed by members of the poly ADP-ribose polymerase (PARP) enzyme family. The removal of PAR chains is mediated by poly ADP-ribose glycohydrolase (PARG), which reverses the process by hydrolyzing ADP-ribose units [3, 4].

Emerging evidence suggests that ADP-ribosylation plays a pivotal role in neurological health, with mutations in genes associated with this pathway linked to epilepsy, autism, schizophrenia and multiple sclerosis [5]. Furthermore, recent studies have implicated ADP-ribosylation in neurodegenerative diseases through its role in protein aggregation. Proteins implicated in diseases such as amyotrophic lateral sclerosis (ALS), frontotemporal degeneration (FTD), and Parkinson’s disease (PD) including TDP-43, FUS, and α-synuclein are PAR binding proteins that are influenced by PARylation, which affects their cellular localization, aggregation, and neurotoxicity [6–11]. Elevated PARylation levels have been observed in motor neurons of ALS patients and in the cerebrospinal fluid of PD patients [9, 10]. Notably, PAR chains, along with several PARPs and PARG isoforms, are associated with cytoplasmic stress granules (SGs) [12].

In vitro studies demonstrate that free PAR chains can initiate the aggregation of critical disease-associated proteins, including TDP-43, FUS, hnRNPA1, and α-synuclein [13].

Recent studies have proposed PARP inhibition as a potential strategy to modulate SG dynamics, protein phase separation and aggregation in ALS [14, 15]. However, these approaches overlook the critical role of PARylation in DNA repair, a process significantly compromised in ALS and other neurodegenerative diseases. PARylation is essential for maintaining genomic stability, as it facilitates the recruitment of repair proteins to sites of DNA damage and stabilizes repair complexes, ensuring efficient resolution of single- and double-strand breaks [16, 17]. This crucial signaling function underscores the risks of PARP inhibition, which could disrupt DNA repair processes.

FUS (Fused in Sarcoma) is an RNA/DNA-binding protein (RDBP) involved in essential cellular processes such as RNA splicing, transport, and stability [18–20] . Mutations in the *FUS* gene are linked to a variety of neurodegenerative disorders, most notably ALS, a progressive disease that leads to the degeneration of motor neurons, resulting in muscle weakness and atrophy [21–23]. ALS is often characterized by the accumulation of misfolded, aggregated proteins, including TDP-43, another RDBP, which can overlap with FUS pathology [24–26]. These protein aggregates disrupt cellular functions and contribute to neuronal toxicity. FUS mutations are also associated with other motor neuron diseases (MNDs), such as FTD, where similar pathologic processes involving RDBP dysfunction occur [21, 27–29]. In both ALS and FTD, there is a shared theme of disrupted RNA metabolism and protein homeostasis, making the study of FUS and its role in disease pathogenesis particularly important. Recent research has focused on the interplay between protein aggregation, DNA repair mechanisms, and the potential therapeutic effects of targeting these pathways [30, 31].

Our group has previously demonstrated specific deficiency in DNA strand break ligation both in nuclear and mitochondrial genome in FUS-associated ALS models [32, 33]. This deficiency correlated with significant accumulation of DNA strand breaks in patient cells expressing FUS mutations, mouse models and in FUS-ALS patient brains. In this context, it is reasonable to perceive that while PARP inhibition may alleviate protein aggregation, it risks exacerbating underlying DNA repair defects, potentially worsening disease pathology.

In this study, we investigated a novel approach to address ALS pathology by expressing a naturally occurring PARG isoform, PARG99, that localizes specifically to the cytoplasm. This isoform targets the protein aggregation component of ALS pathology while preserving nuclear PARylation essential for DNA repair. Our findings demonstrated that while PARP inhibitors had a similar effect on FUS aggregation and SG disassembly, they also resulted in persistent genome instability, as expected, underscoring the critical role of PARylation in maintaining genomic stability. This highlights a clear advantage of our cytosol-specific PARG99 approach over pan-PARP inhibition, as it mitigates protein aggregation without exacerbating DNA repair defects. The PARG99 strategy offers a focused intervention to address neurodegenerative processes associated with ALS without further compromising genomic stability. While our studies in this manuscript focused on FUS toxicity in ALS patient-derived FUS mutant cells, this approach may have broader potential for other neurodegenerative diseases involving PAR-dependent protein aggregation and SG-linked pathology, such as TDP-43 and α-synuclein proteinopathies.

## Materials and Methods Cell lines, and cell culture

Human fibroblasts were grown in Dulbecco’s modified Eagle’s media (DMEM)/F12 media supplemented with 10% fetal bovine serum (FBS), 100U/ml penicillin, 100U/ml streptomycin, 1% MEM non-essential amino acids (GibcoTM), and 1.6% Sodium bicarbonate (Corning). Human embryonic kidney HEK293 (ATCC) cell lines were cultured in DMEM high glucose media supplemented with 10% FBS and 100U/ml penicillin, and 100U/ml streptomycin [32].

### NPSC induction and motor neuron differentiation from iPSC lines

The origin and conversion of healthy control and ALS patient-derived FUS mutant fibroblasts into Human iPSC cells were previously described [34]. iPSC cells were cultured on Geltrex LDEV-free basement membrane matrix coated dishes coated with, with 1X Essential 8media (Cat#A1517001). To derive NPSCs, PSC neural induction media (Gibco A1647801) was used following the manufacturer’s protocol. Approximately 24 hours after sub-plating the iPSCs, the Essential 8 media was replaced with PSC neural induction media, which was maintained for 7 days. The first passage (P0) NPSCs were transferred to Geltrex-coated 6-well plates (Thermo Fisher) and cultured in StemPRO Neural Stem Cell SFM media (Cat# A1050901). Neural induction efficiency was evaluated at the third passage using immunofluorescence staining with the neural stem cell marker Nestin and the pluripotency marker Oct4. Motor neurons were derived from iPSCs obtained from WiCell Research Institute and VIB-KU Leuven, following established protocols with some modifications. iPSC clones were first transferred from a 60-cm dish to a T-25 flask containing neuronal basic media, a 50:50 mixture of Neurobasal and DMEM/F12 media, supplemented with N2 and B27 (without vitamin A). To facilitate cell suspension, iPSC clones were digested with collagenase type IV. The resulting cell spheres were then treated with various inhibitors, including 5 μM ROCK inhibitor (Y-27632), 40 μM TGF-β inhibitor (SB 431524), 0.2 μM bone morphogenetic protein inhibitor (LDN-193189), and 3 μM GSK-3 inhibitor (CHIR99021). This was followed by a 4-day incubation in neuronal basic media containing 0.1 μM retinoic acid (RA) and 500 nM Smoothened Agonist (SAG). Afterward, the spheres were incubated for 2 more days in neuronal basic media supplemented with RA, SAG, 10 ng/ml Brain-Derived Neurotrophic Factor (BDNF), and 10 ng/ml Glial cell-derived neurotrophic factor (GDNF). To dissociate the spheres into single cells, they were exposed to neuronal basic media containing 0.025% trypsin and DNase in a 37°C water bath for 20 minutes. The dissociated cells were then transferred to media containing 1.2 mg/ml ROCK inhibitor to maintain viability. After cell counting, a specific number of cells were seeded onto Laminin-coated dishes or chamber slides (20 μg/ml) and incubated for 5 days in neuronal basic media containing RA, SAG, BDNF, GDNF, and 10 μM DAPT. The media was then switched to one containing BDNF, GDNF, and 20 μM γ-secretase inhibitor (DAPT) for an additional 2 days. For motor neuron maturation, the cells were cultured for more than 7 days in media containing BDNF, GDNF, and 10 ng/ml ciliary neurotrophic factor (CNTF).

### Antibodies and plasmids

The rabbit anti-FUS antibody (Cat# A300–302A) was purchased from Bethyl Laboratories, Inc. The mouse anti-HA (Cat# 2367S), rabbit anti-Phospho Histone H2A.X (Cat# 9718S), and rabbit anti-Poly/Mono-ADP Ribose (E6F6A) antibodies were obtained from Cell Signaling. The mouse anti-G3BP1 (Cat# 13057-2-AP), rabbit anti-TIA1 (Cat# 12133-2-AP), and mouse anti-GAPDH (1E6D9) antibodies were acquired from Protein Tech. Fluorescent secondary antibodies, Alexa Fluor 488 anti-mouse (Cat# A28175) and Alexa Fluor 647 anti-rabbit (Cat# A22287), were sourced from Life Technologies. Antibodies were used at dilutions of 1:1000 for western blotting, 1:500 for immunofluorescence, and 1:100 for PLA.

### Transfection, immunoblotting, and Microscopy

NPSC cells were transfected using Lipofectamine Stem transfection reagent (Cat# STEM00001) (Thermo Fisher), according to the manufacturer’s protocol. Immunoblotting and immunofluorescence were carried out according to standard protocols. For immunoblotting, cell lysates were prepared in 1X RIPA buffer (Millipore) containing a protease inhibitor cocktail (Roche), and proteins were separated by electrophoresis on 4–12% Bis-Tris precast gels (Bio-Rad). After transferring to nitrocellulose membranes, proteins were incubated with primary and secondary antibodies, and chemiluminescence reagents (LI-COR) were used for signal detection, with visualization on a LI-COR Odyssey imaging system. For immunofluorescence, cells grown on chamber slides were fixed with 4% paraformaldehyde for 15 minutes, followed by permeabilization with 0.5% Triton X-100 for 15 minutes. Cells were then incubated with primary antibodies overnight and with fluorescently labeled secondary antibodies for 2 hours. Microscopy was performed using FLUOVIEW FV3000 series of confocal laser scanning microscope.

### Soluble insoluble fractionation

To separate soluble and insoluble protein fractions, cell pellets were first processed for soluble fractions using 1× RIPA buffer containing a protease inhibitor. After high-speed centrifugation, the supernatant was designated as the soluble fraction. The pellet, containing the insoluble protein fraction, resuspended in an equal volume of buffer comprising 2% SDS, 50 mM tris buffer (pH 8.0), and 10% glycerol, followed by sonication. For gel loading, the protein concentration of the soluble fraction was used as a reference for quantifying the insoluble fractions [35].

### Nuclear cytosolic fractionation

Nuclear cytosolic fractionation was performed following protocol published by V V Senichkin et al., 2021. Briefly, cells were suspended in 1 mL of a hypotonic solution containing 0.1% NP-40 and incubated for 3 minutes. The cells were then homogenized using a Potter-Elvehjem homogenizer, performing approximately 20 up-and-down strokes with the pestle. The homogenate was centrifuged at 1,000 rcf for 5 minutes to pellet the nuclei, after one more wash with isotonic buffer the pellet was resuspended in 1XRIPA buffer to get nuclear fraction. The supernatant was subsequently centrifuged again at 15,000 rcf for 3 minutes to remove debris to get cytosolic fraction.

### Long amplicon PCR (LA-PCR)

Genomic DNA was extracted using the Qiagen Blood and Tissue Kit, according to the manufacturer’s instructions. In this study, a 10 kb fragment of the *hPRT* gene (encompassing exons 2–5; accession number J00205) was amplified using LongAmp Taq DNA polymerase (New England Biolabs) with the forward primer 5′-TGGGATTACACGTGTGAACCAACC-3′ and the reverse primer 5′-GCTCTACCCTGTCCTCTACCGTCC-3′. As a control, a shorter 250 bp fragment of the *hPRT* gene was amplified using the forward primer 5′- TGCTCGAGATGTGATGAAGG-3′ and the reverse primer 5′-CTGCATTGTTTTGCCAGTGT- 3′ [33, 36]. Following amplification, the PCR products were analyzed by two independent methods: agarose gel electrophoresis and the Quant-iT PicoGreen DNA quantification assay (Thermo Fisher).

### CellTiter-Glo 2.0 viability assay

NSC cells were seeded onto white opaque 96-well plate and exposed to 100ng/ml GO for 1 h. After treatment, media was replaced with culture media and cells were allowed to grow for 24 hours. The plate and its contents were equilibrated to room temperature for approximately 30 minutes. An equal volume of CellTiter-Glo 2.0 reagent (Promega-G9242) was then added to match the cell culture medium in each well [37]. The mixture was gently agitated on an orbital shaker for 2 minutes to facilitate cell lysis, followed by a 10 minute incubation at room temperature to stabilize the luminescent signal. Luminescence was subsequently measured using a Tecan microplate reader. This luminescent signal reflects ATP levels in the sample, serving as a marker for viable, metabolically active cells.

### Statistical analyses

Each dataset presented represents at least three independent experiments. Statistical analysis was conducted using GraphPad Prism software, and significant differences were assessed using Student’s t-test. A p-value of less than 0.05 was considered statistically significant.

## Results

### FUS pathology alters SGs dynamics and PARP inhibitor treatment enhances their resolution

To investigate the impact of FUS mutations on SG dynamics, we utilized two cell lines derived from ALS patients harboring established, pathogenic FUS mutations P525L and R521H. Patient fibroblasts (Supplementary Fig. 1) and motor neurons differentiated from FUS-P525L iPSCs (Fig. 1) were treated with sodium arsenite (500μM/30 min) to induce SG formation (Fig.1A). Treated cells were subjected to immunofluorescence (IF) staining with antibodies against FUS, and G3BP1, a marker of SGs [38] (Fig. 1B and supplementary Fig. 1). Both WT and mutant cells formed SGs, however, mutant cells exhibited a greater number of SGs with increased FUS localization.

**Fig. 1:**
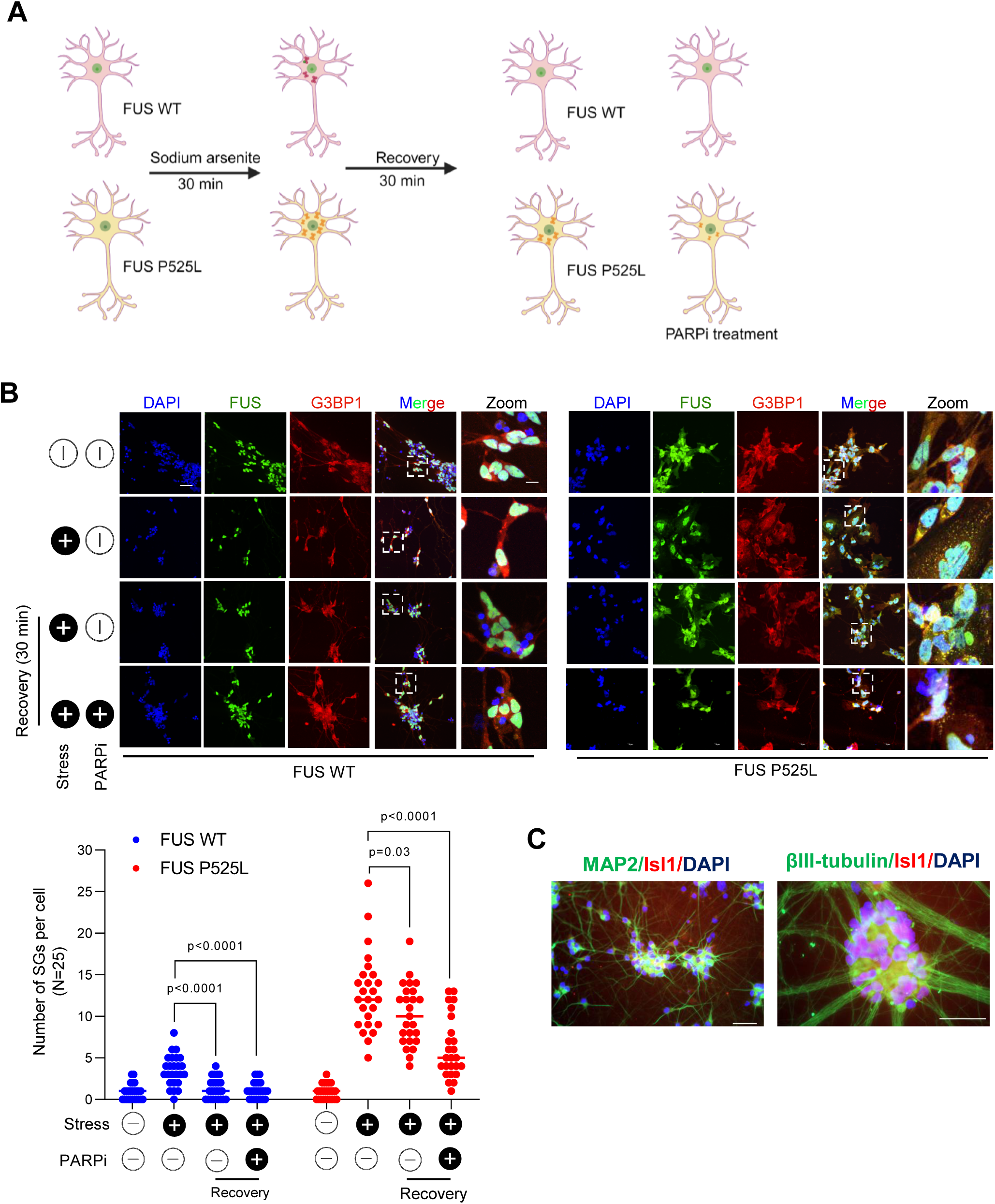
FUS pathology increases stress granule (SG) formation and delays SG resolution, while PARP inhibitor (PARPi) treatment enhances SG resolution. A. Schematic of experimental design and PARPi treatment plan. B. Immunofluorescence (IF) images of control (FUS WT) and patient (FUS P525L) iPSC derived motor neurons with Mock (PBS), sodium arsenite for 30 minutes, and 30 minutes recovery after sodium arsenite treatment in presence and absence of PARP inhibitor (Veliparib). Cells were stained with antibodies against FUS in green, G3BP1 in red and nucleus counter stained with DAPI. Scale bar = 10μM for full images; scale bar = 3μM for zoomed images. Quantification data represented as mean ± s.e.m derived from three independent experiments with SG quantification performed on 25 cells per condition. Statistical analyses were performed using a two-sided Student’s-t test (Graphpad prism software). C. IF staining of motor neuron markers: representative images demonstrate motor neurons positively for Isl-1, MAP2, or βIII-tubulin, indicating an approximate differentiation efficiency of 80%. Scale bar = 50µm.

We then performed recovery experiments to evaluate SG resolution by allowing the cells to recover is fresh media for 30 minutes, following sodium arsenite treatment for 30 minutes. The WT cells recovered more efficiently from stress compared to cells harboring FUS mutations (Fig. 1B). Motor neuron differentiation from iPSCs resulted in 80% differentiation efficiency, the differentiation is validated by expression of neuronal marker MAP2 and motor neuron marker Isl-1 (Fig. 1C)

To confirm that the observed changes in SG dynamics were due to FUS mutations, we used isogenic control lines in which the FUS mutations were corrected by CRISPR-Cas9 knock-in [34]. These corrected lines exhibited comparable SG formation and resolution in response to stress, closely resembling WT cells and showing distinct differences from the mutant cells (Supplementary Fig. 3). These results suggest that the observed changes in SG dynamics can be attributed specifically to FUS mutations.

To further explore the effects of PARP inhibition (PARPi) on SGs formation and resolution, cells were pre-treated with veliparib (10μM) for 30 minutes, before sodium arsenite exposure. Following arsenite treatment, cells were allowed to recover in fresh media containing PARPi for 30 minutes (Fig. 1A). This approach was tested in both patient-derived fibroblasts and iPSC derived motor neurons (Fig. 1B and Supplementary Fig. 2). In both cases, PARPi treatment led to a significant reduction in SGs number and size during recovery.

Importantly, the ability of PARP inhibition to facilitate SG resolution even in mutant cells underscore the involvement of PARylation activity of PARPs in SG dynamics in motor neurons. This also raises the question about the functional versus pathological significance of PARylation in SG dynamics and the cellular stress response.

### Inhibition of PARPs enhances solubility of RDBPs TDP-43 and FUS but exacerbates genome instability

Cytoplasmic mislocalization of FUS is often considered a precursor to its pathology in ALS, although direct evidence linking its localization to solubility remains limited [39]. Here, we compared the protein solubility between wild type (WT) FUS and patient derived P525L FUS mutant fibroblasts cells using cellular fractionation. Soluble proteins were extracted using 1X RIPA buffer, while insoluble proteins were isolated using a 2% SDS buffer [35]. Immunoblotting analysis revealed a significant increase in insoluble FUS in P525L mutant cells compared to WT cells (Fig. 2). Notably, no significant changes were observed in insoluble TDP-43 levels under unstressed conditions (Fig. 2). However, upon sodium arsenite treatment, we observed a shift in proteins, such as SG-associated proteins G3BP1, TIA1, and ALS-associated RDBPs TDP-43, and FUS, from soluble to the insoluble fraction. This shift was accompanied by an increase in ADPR levels, which were predominantly enriched in the insoluble fraction, further emphasizing the effect of stress on protein solubility. GAPDH levels in the soluble fraction were used as loading control.

**Fig. 2:**
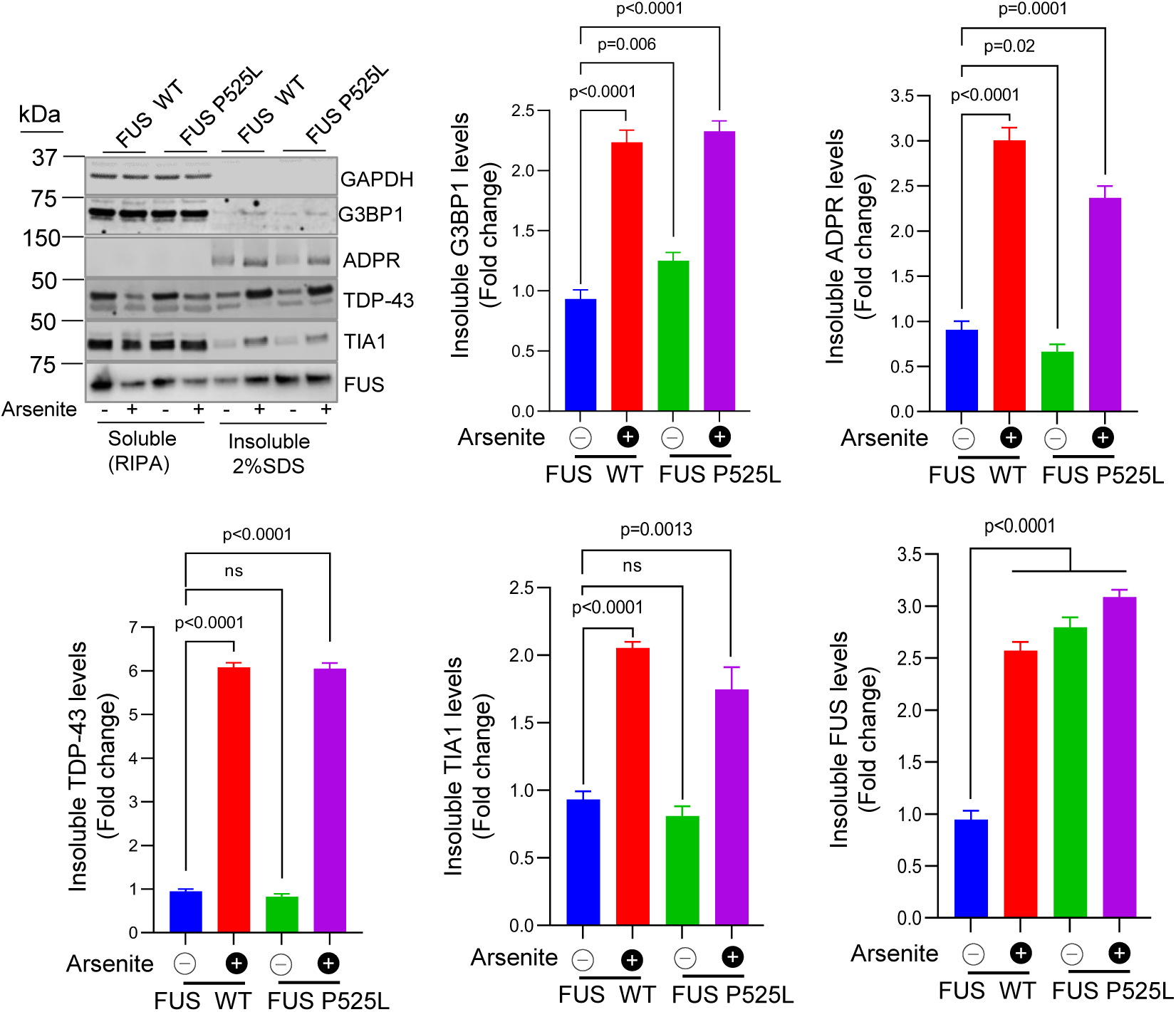
FUS pathology and sodium arsenite treatment alter protein solubility. Immunoblots (IB) of soluble (1X RIPA) and insoluble (2% SDS) protein fractions from control (FUSWT) and patient derived (FUS P525L) fibroblasts following sodium arsenite treatment (500μg/30min). Insoluble protein levels were quantified, and data represented as mean ± s.e.m from three independent experiments. Statistical analyses were performed using two-sided Student’s-t test (Graphpad prism software).

Next, we examined the impact of PARPi on protein solubility following sodium arsenite treatment. NPSC cells treated with arsenite in the presence or absence of the PARPi Veliparib showed a significant reduction in the insoluble protein levels (Fig. 3). We also analyzed the solubility of FUS, TDP-43, G3BP1 and TIA1 as before. The reduction in these proteins in insoluble fractions corresponded to an increase in their soluble forms after PARPi treatment (Fig. 3). These findings suggest that PARP inhibition reduces the formation of insoluble protein aggregates.

**Fig. 3:**
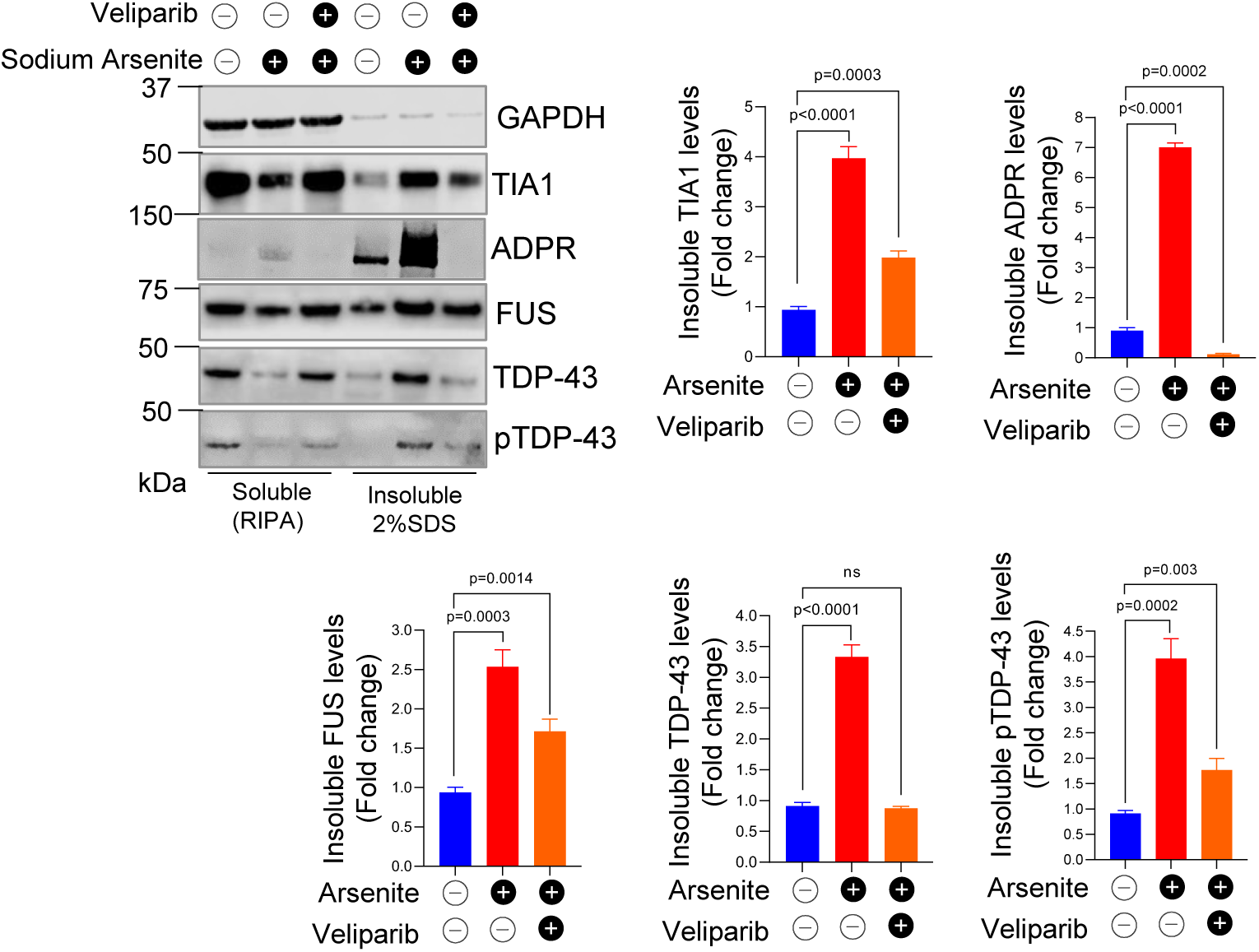
PARP inhibitor (Veliparib) reduces protein insolubility after sodium arsenite treatment. Immunoblots (IB) of soluble (1X RIPA) and insoluble (2% SDS) protein fractions showing the soluble levels of indicated proteins in FUS WT neuronal progenitor stem cells (NPSC’s) treated with sodium arsenite, with or without Veliparib. Quantification of insoluble protein levels presented as mean ± s.e.m from three independent experiments. Statistical analyses were performed using two-sided Student’s-t test (Graphpad prism software).

A common feature of both FUS and TDP-43 associated ALS pathology is a defective DNA repair response, a hallmark also shared by many neurodegenerative diseases [33, 40, 41]. While PARPi has shown potential to improve protein solubility, we hypothesize that their longterm application as a therapeutic approach for ALS may be limited due to their detrimental impact on DNA repair pathways. Efficient DNA repair is critical for maintaining genomic stability, and impairments in this process could worsen the underlying pathology in ALS. To investigate this, we utilized iPSC-derived motor neurons from both FUS WT and P525L mutant cells. These cells were treated with glucose oxidase (GO), a DNA-damaging agent, for 1 hr, followed by a 6-hr recovery in the presence or absence of PARPi veliparib (Fig. 4A). DNA damage and repair efficiency were assessed by fixing cells at three key time points: before GO treatment, immediately after treatment, and following the recovery period. Immunofluorescence evaluation of γH2AX foci formation was used as a marker for DNA damage, with efficient DNA repair indicated by a decrease in foci during recovery (Fig. 4B).

**Fig. 4:**
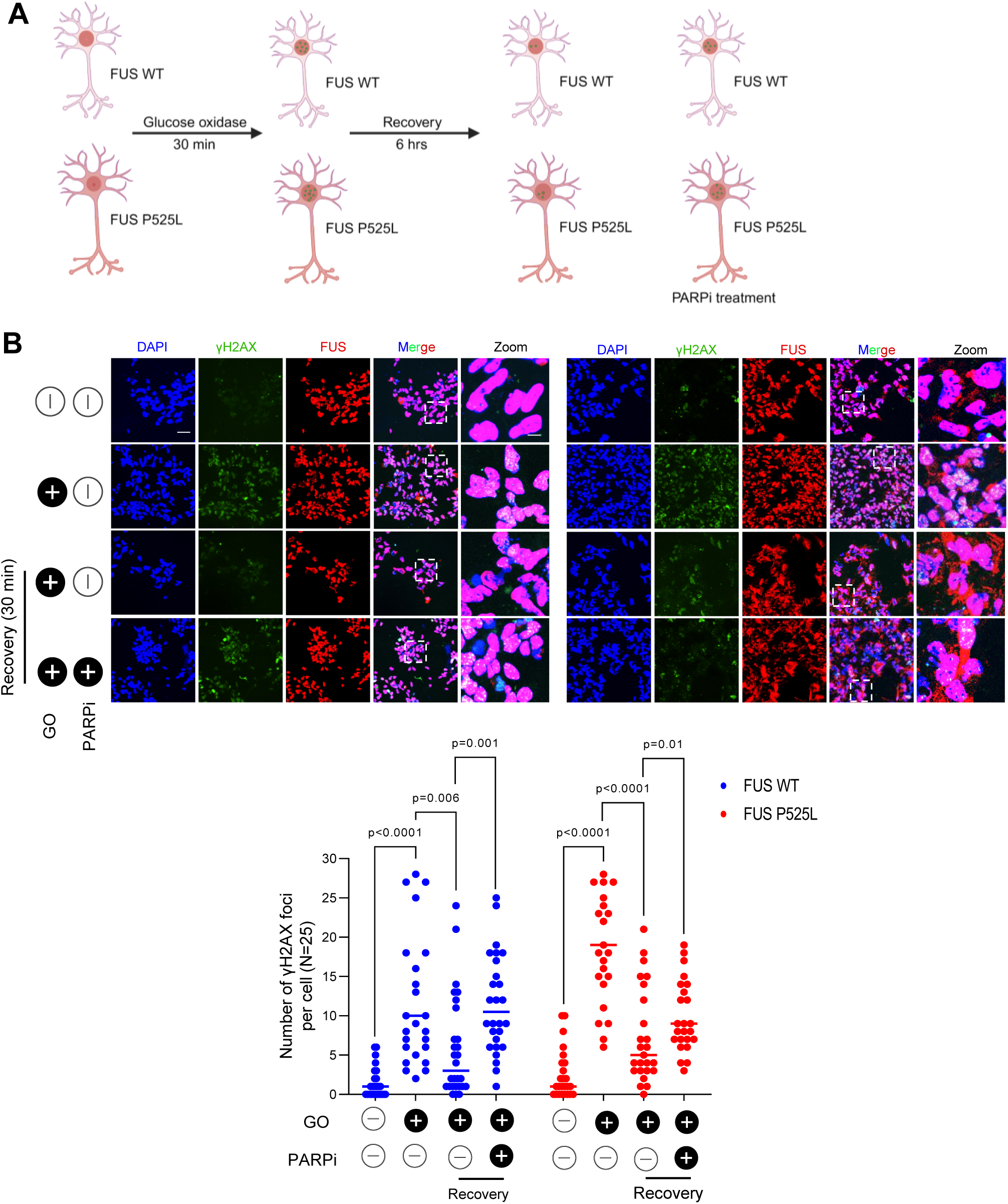
PARP inhibitor treatment causes delayed γH2AX foci resolution after glucose oxidase (GO) treatment. A. Schematic of the experimental design. B. Immunofluorescence (IF) images of control (FUS WT) and patient (FUS P525L) iPSC derived motor neurons with Mock (PBS), GO (100 ng/ml) for 30 min, followed by 6 hrs of recovery in the presence or absence of PARP inhibitor (Veliparib). Cells were stained with antibodies against FUS (red), γH2AX (green) and nuclei counter stained with DAPI (blue). Scale bar = 20 μM for zfull images; scale bar = 3 μM for zoomed images, Quantification data represented as mean ± s.e.m from three independent experiments, with quantification of γH2AX foci from 25 cells per condition. Statistical analyses were performed using two-sided Student’s- t test (Graphpad prism software).

Our findings revealed that although there were some differences in the degree of clearance related to FUS pathology and DNA repair defects, both FUS WT and P525L mutant cells were able to recover from DNA damage to a significant extent, as evidenced by the reduction in the number of γH2AX foci during the recovery phase (Fig. 4B). However, cells treated with PARPi exhibited a marked failure to resolve γH2AX foci, with high levels persisting even after the recovery period (Fig. 4B). These results suggest that while PARPi might transiently improve RDBP protein solubility, its negative impact on DNA repair capacity, which is already disrupted in ALS pathology, could worsen the long term pathological outcomes, underscoring the need for caution when considering PARPi as a therapeutic strategy.

### Subcellular PARG isoform expressions and their effect on RDBP solubility and DNA damage response

Previous studies have shown that PARG (full length) expression produces similar outcomes in ALS pathology to PARPi in the context of protein solubility [14]. However, this strategy has similar limitations as PARPi, due to the DNA repair defects associated with total cellular dePARylation [14, 42]. We hypothesize that a better approach to address this issue would involve targeted dePARylation in cytoplasm, without affecting the nuclear PARylation, which is important for DNA repair. We tested this hypothesis by utilizing subcellular compartment specific PARG isoforms, taking advantage of naturally occurring PARG variants that predominantly localize to cytosol [4]. We used two PARG variants, PARG111, a full-length protein with both N-terminal nuclear localization signal (NLS) and nuclear export signal (NES), which expresses in both the nucleus and cytoplasm; and PARG99 which lacks the NLS and is exclusively localized to the cytoplasm (Fig. 5A). These constructs were generated from cDNA of WT NPSC cells, and we confirmed the sequence of the open reading frame (ORF) and validated their expression through immunoblotting, followed by immunofluorescence to confirm their cellular localization (Figs. 5B and 5C).

**Fig. 5:**
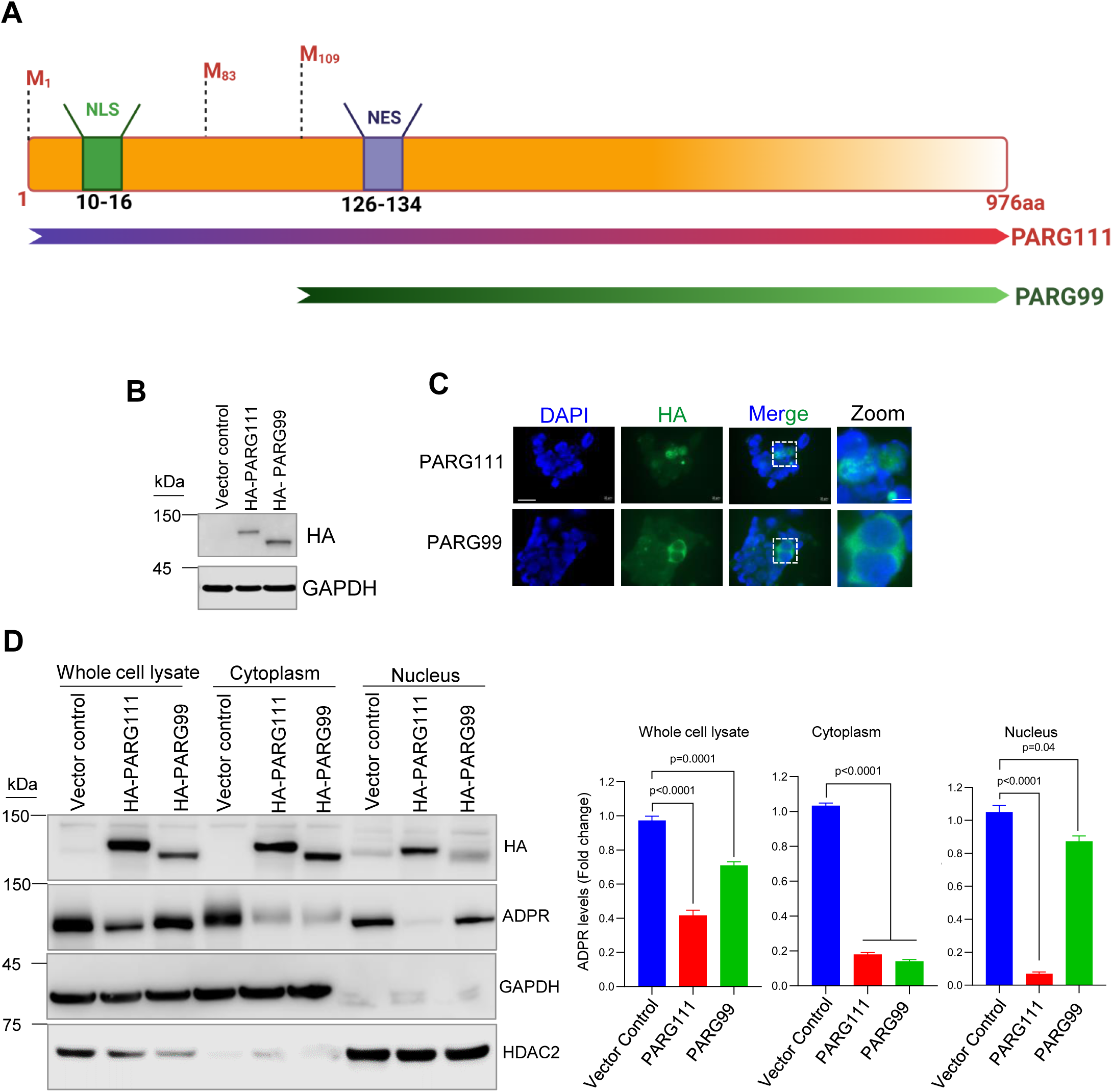
PARG isoforms and their subcellular localization. A. Schematic diagram depicting PARG isoforms. Full length PARG (PARG111) and cytosolic PARG (PARG99) are the products of a single gene and produced naturally through alternative splicing. Translational start sites, nuclear localization signal (NLS), and nuclear export signal (NES) are indicated. B. Immunoblots (IB) showing HA-PARG isoform expression in HEK293 cells. C. Immunofluorescence (IF) images showing HA-PARG localization in HEK293 cells, HA stained in green, and nucleus was counter stained with DAPI (blue). Scale bar = 10 μM for full images; scale bar = 1 μM for zoomed images. D. IB of cellular fractionation in PARG isoform-expressing HEK293 cells, showing their effects on ADP ribosylation (ADPR). Quantitation of ADPR levels in each cellular compartment is shown. Data are presented as mean ± s.e.m. from three individual experiments. Statistical analyses were performed using two-sided Student’s-t test (Graphpad prism software).

Next, we examined the effect of PARG variant expressions on cellular ADPR levels. We transfected the plasmids into HEK293 cells and performed cellular fractionation to isolate whole cell lysates, nuclear fractions, and cytoplasmic fractions. The purity of the fractions was confirmed by the absence of GAPDH in nuclear fractions and the lack of HDAC2 in the cytoplasmic fractions (Fig. 5D). We observed that PARG111 expression resulted in a marked decrease in ADPR levels across all three fractions, as expected. However, PARG99 expression led to a moderate decrease in ADPR in the whole cell lysate and no significant decrease in nuclear fractions, suggesting that nuclear PARylation was unaffected after cytosolic PARG99 expression. In the cytoplasm, where the PARG99 is exclusively localized, there was a significant decrease in ADPR levels, supporting the idea of subcellular compartment-specific PARylation inhibition (Fig. 5D).

We then tested the effects of PARG variant expressions on RDBP protein solubility and SG formation, using FUS P525L mutant neuronal progenitor stem cells (NPSC) cells. PARG111 and PARG99 plasmids were transfected into these cells, followed by treatment with sodium arsenite for 30 minutes. Soluble and insoluble fractions were prepared, and immunoblotting was performed to analyze protein levels. We observed a decrease in ADPR induction in PARG- expressing cells, which corelated with a reduction in insoluble forms of FUS, TDP-43, G3BP1 and TIA1 (Fig. 6A). This suggests that similar to PARPi treatment, expression of PARG isoforms improve protein solubility.

**Fig. 6:**
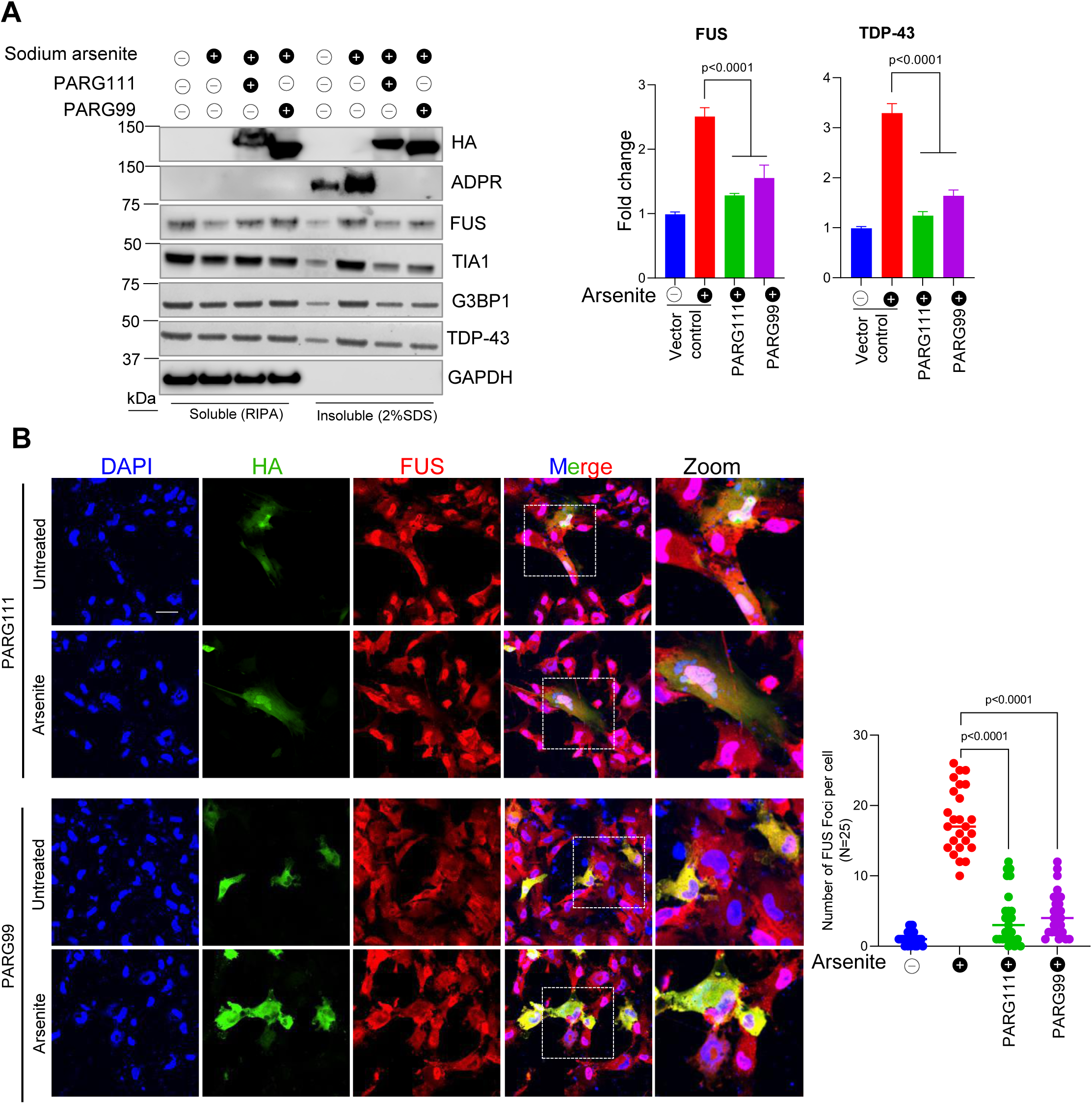
PARG expression increases protein solubility and prevents FUS foci formation after sodium arsenite treatment. A. Immunoblot (IB) of endogenous GAPDH, FLAG, G3BP1, ADPR, TDP-43, TIA1 and FUS in FUS WT NPSC’s. Quantitation histogram represents FUS and TDP-43 levels in the insoluble fractions under the indicated treatment conditions. B. Immunofluorescence (IF) images of FUS P525L iPSC-derived motor neurons transfected with full length (HA-PARG111) and cytoplasmic HA-PARG99. HA-PARG is detected in green, FUS in red using appropriate antibodies and nuclei counter stained with DAPI (blue). Scale bar = 10 μM for full images; scale bar = 3 μM for zoomed images. Data are presented as mean ± s.e.m. from three individual experiments. Statistical analyses were performed using two-sided Student’s-t test (Graphpad prism software).

Next, we investigated the effect of PARG variants on SG formation after sodium arsenite treatment. To specifically examine FUS aggregation, we used FUS P525L mutant motor neurons, transfected them with both PARG variants. After transfection, the cells were treated with sodium arsenite, and experimental groups before and after treatment were fixed for immunofluorescence, and stained for antibodies against FUS and HA, a tag expressed along with PARG variants (Fig. 6B). The expression pattern of PARG variants was consistent with our previous observation in cellular fractionation experiment PARG111 was expressed in both nuclear and cytoplasm, while PARG99 localized exclusively to the cytoplasm. Quantification of SGs, revealed by immunofluorescence staining for G3BP1, indicated a significant decrease in cells expressing PARG variants, in comparison to the nearby non-transfected cells in the same microscopic field. Our findings suggest that even low to moderate levels of PARG expression are sufficient to effectively resolve SGs, even under conditions of cellular stress.

These results highlight that both full length and cytoplasmic-specific PARG variants can effectively mitigate RDBP protein aggregation and SG accumulation, similar to PARPi, suggesting that compartmentalized dePARylation may be a promising strategy for addressing ALS-associated protein pathology.

### Effects of PARPi and PARG variant expressions on DNA repair and cell survival

The potential of targeted PARG99 expression to alleviate FUS toxicity, depends on its ability to demonstrate minimal impact on DNA repair compared to PARPi and PARG111. To address this, we employed two distinct methods: long amplification PCR (LA-PCR) and γH2AX foci analyses.

First, we performed LA-PCR in NPSC cells to evaluate DNA damage accumulation defects associated with PARylation inhibition. Using the HPRT gene, we amplified a ∼10 kb DNA fragment alongside a 250 bp short DNA fragment as a control. The results revealed a significant reduction in DNA repair capacity in cells treated with PARPi or expressing PARG111, which correlated with nuclear ADPR inhibition. In contrast, PARG99 exhibited repair efficiency similar to that of control cells (Fig. 7A).

**Fig. 7:**
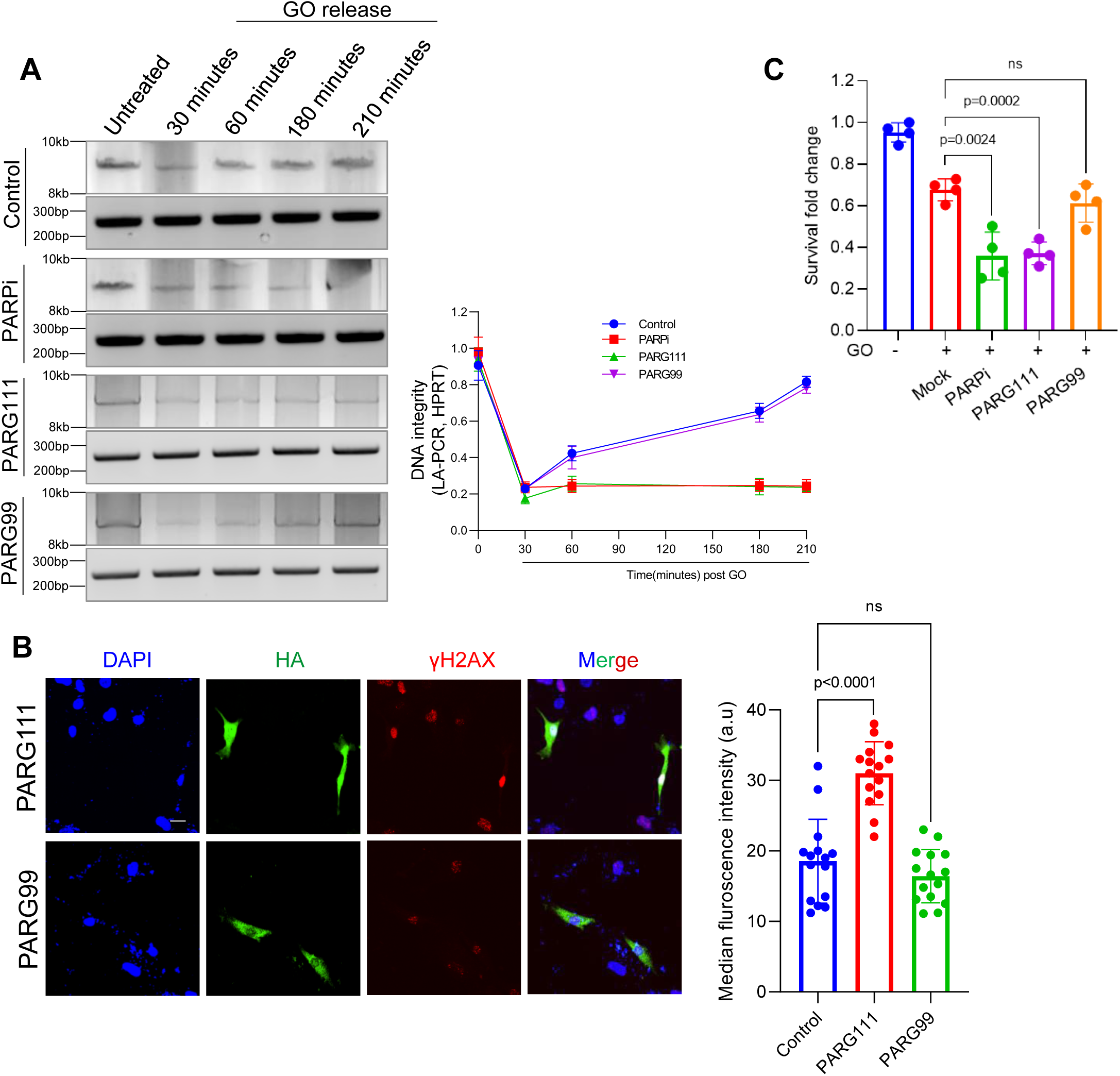
Impact of PARG variant expression and PARPi treatment on DNA repair and cell survival. A. Long amplification PCR (LA-PCR) was performed to assess the genomic integrity after glucose oxidase (GO; 100ng/60 min) treatment in PARG variant-transfected and PARPi-treated NPSC cells. A ∼10 kbp nuclear DNA fragment within the HPRT gene was amplified and resolved on a 1% agarose gel, alongside a 200 bp control PCR product. Amplified PCR products were quantified using PicoGreen fluorescence. B. Immunofluorescence (IF) images of FUS P525L iPSC-derived motor neurons transfected with HA-PARG full length (HA-PARG111) or cytoplasmic (HA-PARG99) variant. HA-PARG shown in green, γH2AX in red using appropriate antibodies and nucleus counter stained with DAPI (blue). Scale bar = 10μM. C. Viability assay using the Cell Titer-Glo kit to assess NPSC cell survival after GO treatment in PARG variant-expressing and PARPi-treated cells. Data are presented as mean ± s.e.m. from three individual experiments. Statistical analyses were performed by two-sided Student’s-t test (Graphpad prism software).

To further validate the DNA repair efficiency between the PARG variants, we compared γH2AX foci formation in PARG111 and PARG99 expressing cells after treatment with the genotoxic GO, followed by a (180 min) recovery period. This condition mirrors the conditions used for PARPi treatment in Fig. 7A. We observed reduced foci formation in PARG99-expressing cells, comparable to the non-transfected control cells. However, PARG111-expressing cells showed a significant increase in foci formation (Fig. 7B), indicating a higher level of DNA damage in these cells. This suggests that PARG99 does not impair DNA repair, unlike PARG111 or PARPi treatment.

Finally, we assessed cell survival under genotoxic stress, using Cell Titer-Glo assay. Cells transfected with PARG variants and treated with PARPi were exposed to GO for 1 hr, followed by a 6 hr recovery in fresh media. Cells expressing PARG99 showed cell survival rates comparable to control cells, whereas PARG111 and PARPi treatment resulted in significant survival defects after GO treatment (Fig. 7C). These results suggest that PARG99 preserves DNA repair efficiency and promotes cell survival under genotoxic stress, offering a better therapeutic alternative than PARG111 and PARPi.

Taken together, our findings corroborate previous studies highlighting the role of PARPi in regulating SG dynamics, while also providing new insights into its broader impact on RDBP protein solubility. However, we also uncovered key limitations associated with PARPi treatment in the context of FUS-ALS, particularly its detrimental effects on DNA repair mechanisms and overall cell survival. These negative consequences underscore the challenges of using PARPi in therapeutic contexts where maintaining faithful DNA repair is crucial. To overcome these limitations, we introduced an novel approach that targets protein aggregation, a hallmark of ALS pathology, through cellular compartment-specific inhibition of PARylation. By leveraging the expression of PARG variants, which localize to specific cellular compartments, we demonstrated how cytosol-specific PARylation inhibition can effectively reduce protein aggregation without interfering with essential nuclear processes like DNA repair (Fig. 8). This approach holds promise as a potential avenue to protect against protein misfolding and aggregation, while preserving critical cellular processes such as DNA repair and cell survival. Thus, this strategy could lead to the development of new treatments for neurodegenerative diseases like ALS.

**Fig. 8:**
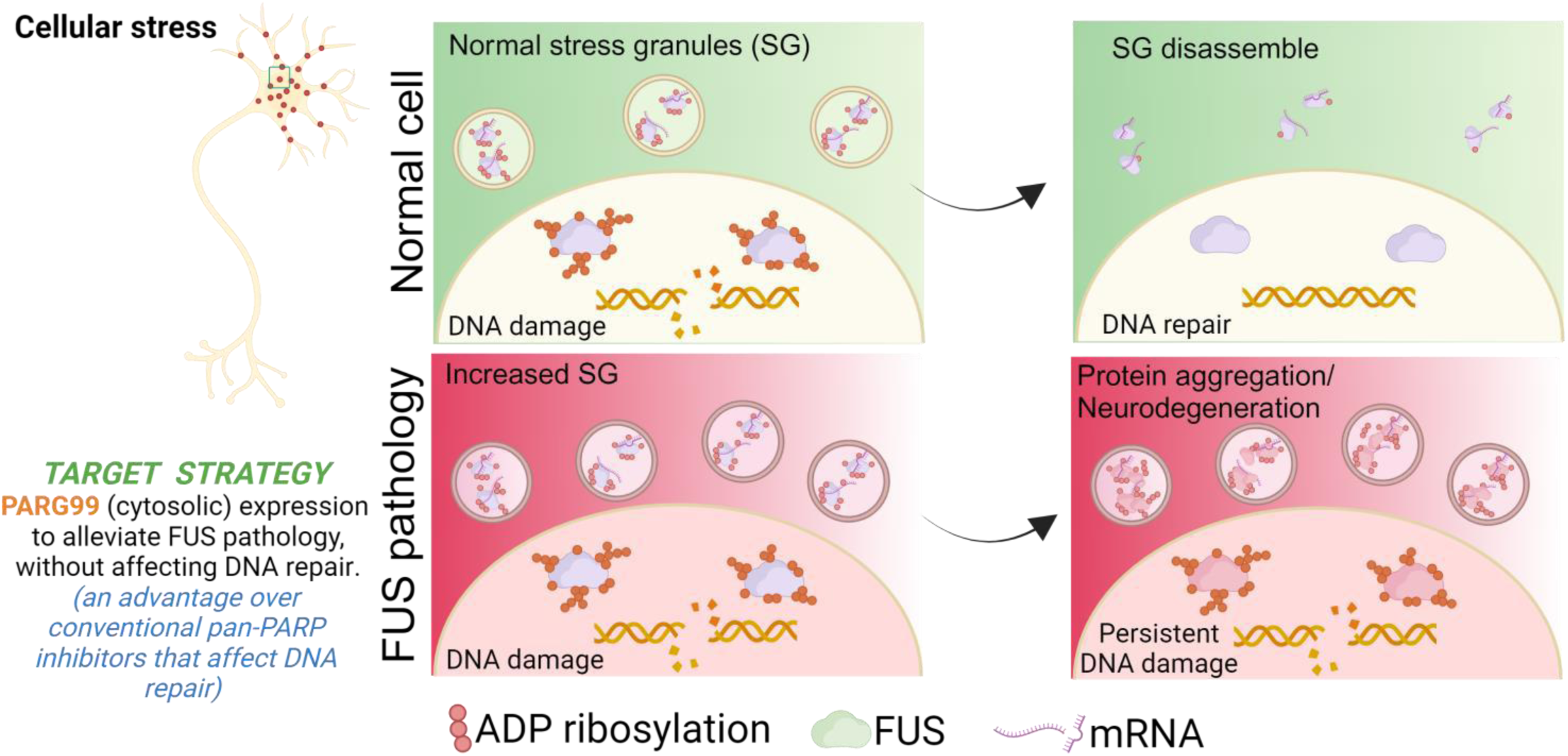
A schematic model summarizing the potential of PARG99 expression in mitigating **FUS pathology-associated protein aggregation.** ADP ribosylation is an important post-translational modification often dysregulated in FUS and other ALS-related pathologies. Conventional approaches utilizing PARP inhibitors (PARPi) can be detrimental to cells due to their adverse effects on DNA repair and other essential cellular functions. In contrast, PARG99 can be expressed in a subcellular compartment-specific manner, reducing protein aggregation without affecting DNA repair, offering a promising avenue for treating RNA/DNA binding protein pathology-associated neurodegenerative diseases.

## Discussion

SGs are dynamic, membrane-less organelles that are transiently formed in response to cellular stress, through liquid-liquid phase separation [43]. While SG formation is a protective stress response mechanism, their dysregulation and persistence are increasingly linked to neurodegenerative diseases, including ALS, frontotemporal dementia (FTD) and Alzheimer’s disease [44, 45]. Transient SGs are typically formed in cultured cells with 30-60 minutes of exposure to stress, to protect critical cellular RNA components. However, chronic stress in age-associated neurodegenerative diseases likely lead to persistent accumulation of unresolved SGs that are often linked to aggregation of disease-associated proteins [46, 47].

FUS localization to SGs has previously been shown in cellular models under overexpression and stress conditions [48–50]. In this study, we employed ALS patient-derived fibroblasts, iPSCs, and differentiated motor neurons with FUS mutations to evaluate SG dynamics, protein solubility, and DNA damage repair. Consistent with prior reports, our study confirms that familial ALS-associated FUS mutations promote enhanced SG formation and delayed resolution under stress conditions, contributing to the pathology of ALS. Sodium arsenite-induced stress led to increased insolubility of key SG marker proteins (e.g., G3BP1, TIA1) and ALS-associated proteins FUS, TDP-43. Notably, the FUS P525L mutation, which is associated with severe juvenile ALS (jALS) [51], exhibited persistent protein insolubility even in the absence of stress, underscoring its severe pathogenicity.

Given the established role of PARylation in SG dynamics and neurodegeneration-associated protein aggregation, pharmacological PARP inhibitors (PARPi) have been proposed as a potential therapeutic strategy [8, 52] . The activation of PARP1 in ALS patient spinal cords and the role of PAR in FUS aggregation [14], further support the concept of PARP inhibition as a target in FUS-ALS. However, our findings highlight significant limitations of this approach. While PARPi effectively modulated SG dynamics and protein aggregation in ALS patient-derived motor neurons, it also exacerbated DNA repair defects, as evidenced by persistent γH2AX foci under chronic stress conditions. This observation aligns with the critical role of PARylation in genomic stability, particularly its facilitation of DNA repair through the recruitment and stabilization of repair complexes [16, 17]. Our previous research identified a functional interaction between FUS and PARP1, which plays a crucial role in two key processes: FUS recruitment to DNA damage sites and FUS-mediated assembly of the DNA break-sealing XRCC1-DNA ligase III complex at break sites [33]. Both processes are dependent on the PARylation activity of PARP1. In this context, global PARP inhibition is anticipated to exert long-term adverse effects on genome stability, as experimentally corroborated in this study.

To address this challenge, we explored a cytoplasm-specific approach using the naturally occurring PARG splice variant, PARG99. PARG, a key enzyme in PAR metabolism, has been linked to neurodegeneration with conflicting outcome, ranging from loss of PARG leading to neurodegeneration to PARG inhibition being considered as a therapeutic strategy [53, 54]. Previous studies suggest that PARG expression has a similar effect on SG dynamics as PARPi but may induce DNA repair defects by inhibiting nuclear PARP activity [14].

Our results demonstrate that unlike full-length PARG (PARG111) or PARPi, PARG99, which predominantly localizes to the cytoplasm, selectively targeted cytoplasmic PARylation without impairing nuclear DNA repair. Notably, both PARG111 and the cytoplasmic PARG99 variants showed comparable effects on SG dynamics and protein solubility following sodium arsenite treatment. However, in the context of DNA repair after GO treatment, PARPi and PARG111 delayed γH2AX foci resolution, whereas PARG99 did not, as evidenced by lower γH2AX foci levels. This finding was further corroborated by long amplicon (LA)-PCR, which demonstrated that PARG99, unlike PARPi and PARG111, enabled DNA repair recovery comparable to control cells. Finally, cell survival assays following genotoxic stress, revealed reduced survival with PARPi and PARG111, while PARG99 maintained survival rates comparable to control cells.

Thus, functional assays revealed that PARG99 preserved DNA integrity, and improved cell survival under genotoxic stress, supporting its potential as a safer alternative to PARP inhibition.

Overall, our findings established two key insights: (1) PARylation is integral to SG dynamics and protein aggregation in ALS pathology, and (2) the cytoplasm-specific targeting of PARylation via PARG99 mitigates protein aggregation while preserving DNA repair, addressing a major limitation of PARP inhibitors. Importantly, these results highlight the potential of PARG99 in alleviating FUS toxicity in ALS and other neurodegenerative diseases characterized by PARylation-associated pathology. Future studies are warranted to explore the broader applicability of PARG99, particularly in diseases involving TDP-43 and α-synuclein proteinopathies. The cytoplasm-specific targeting of PARylation represents a promising avenue for developing targeted interventions that address neurodegenerative mechanisms without compromising genomic stability.

## Acknowledgements

This research is primarily funded by the National Institute of Neurological Disorders and Stroke (NINDS) and the National Institute on Aging of the National Institutes of Health (NIH) under award number RF1NS112719 to M.L.H. Additional support for research in the Hegde laboratory is provided by NIH awards R01NS088645 and R01NS094535, as well as by the Sherman Foundation Parkinson’s Disease Research Challenge Fund and internal funding from the Houston Methodist Research Institute. M.L.H. also acknowledges the support of Everett E. and Randee K. Bernal through the Centennial Endowed Chair of the Neurological Institute. The authors extend their gratitude to the members of the Hegde laboratory for their contributions, and to Drs. Anna Dodson and Gillian Hamilton and at the Houston Methodist Research Institute (Houston, TX) for their editorial assistance and financial support of Walter’s educational initiative.

## Author Contributions

M.K. co-designed and performed experiments with assistance from V.H.M., and J.M. M.K. conducted data analysis with interpretation and co-wrote the manuscript. M.L.H. designed and supervised the study and prepared the final manuscript. All authors participated in the discussion and provided valuable feedback on the manuscript.

## Data availability statement

Original, raw data from the experiments conducted in this study can be provided upon reasonable request to the corresponding authors.

