## Supplementary material for "Selective Inhibition of Cytosolic PARylation via PARG99: A Targeted Approach for Mitigating FUS-associated Neurodegeneration"

**Supplementary Material** accompanying this paper includes three Figures and one table.

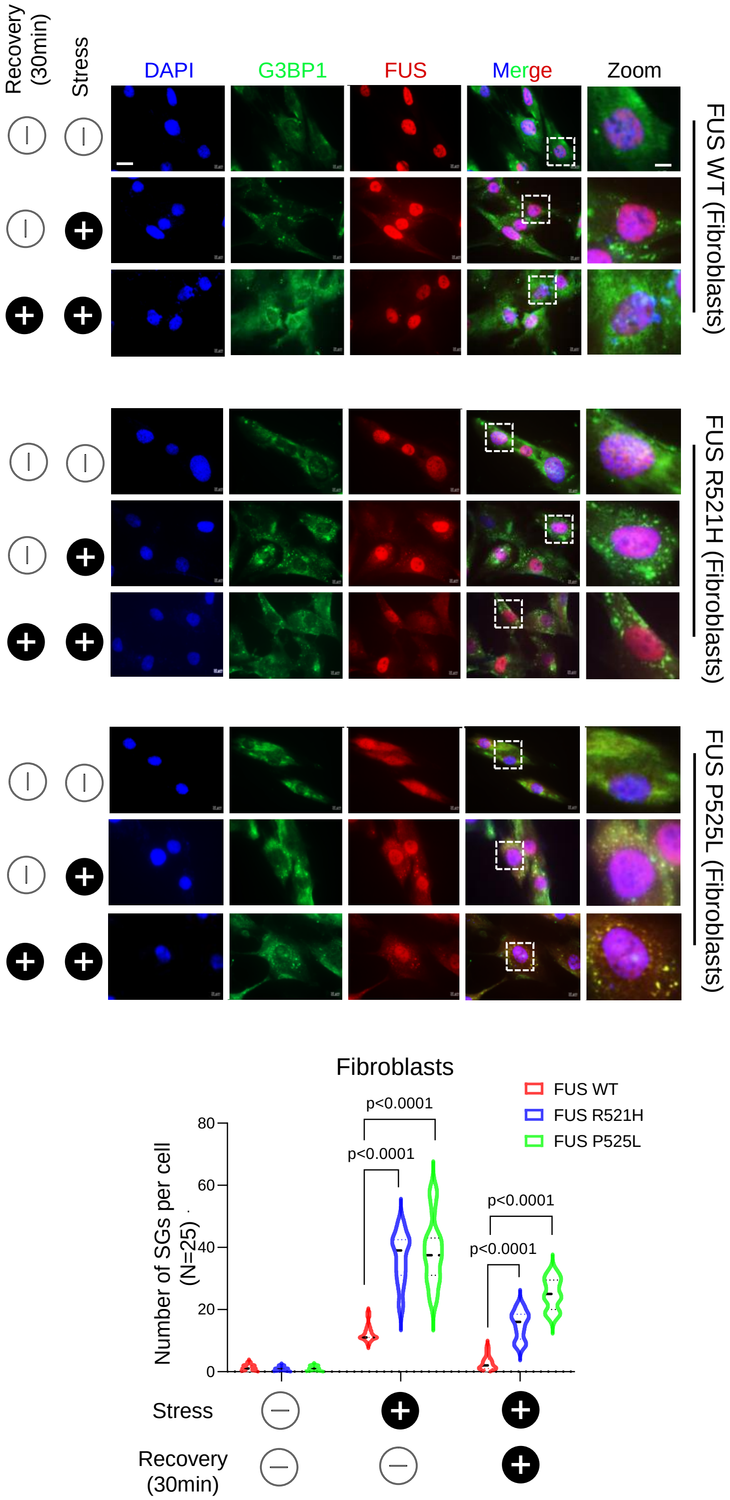


**Supplementary Fig1: FUS pathology causes increased SGs number and delays SGs resolution.** Immunofluorescence images of control and patient derived fibroblasts containing FUS R521H and P525L mutations were treated with PBS, Sodium arsenite for 30 minutes, and 30 minutes recovery after sodium arsenite treatment. Stained with G3BP1 in green, FUS in red and nucleus counter stained with DAPI. Scale bar =20μM for zoomed image scale bar=3 μM, Quantification data represented as mean ± s.e.m derived from three independent experiments, quantification of SGs was derived from 25 cells. All statistical analysis were performed by two-sided student's-t test using graph pad prism software.

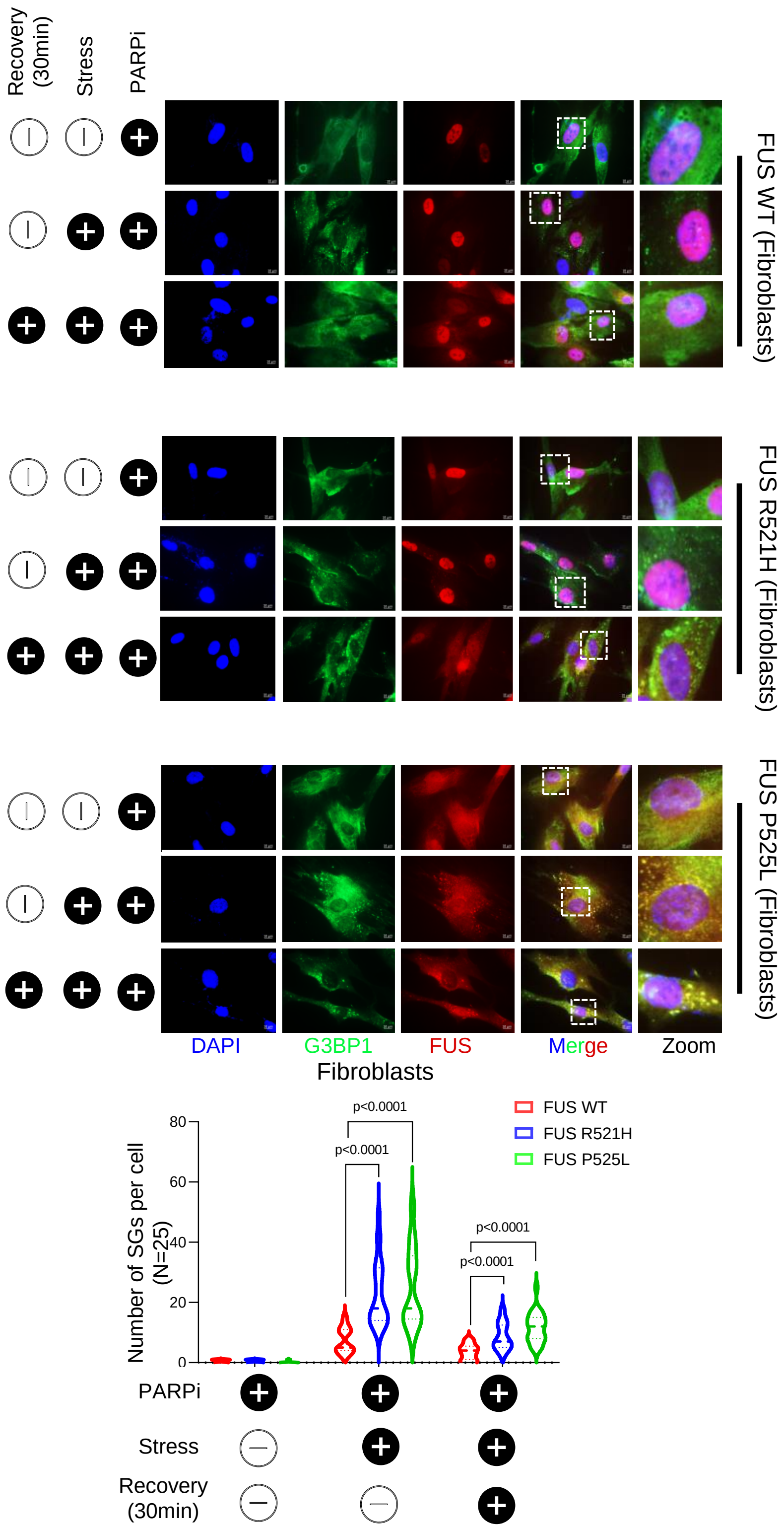


**Supplementary Fig2: PARP inhibitor (PARPi) treatment reduces SGs number and enhances resolution.** Immunofluorescence images of control and patient derived fibroblasts and containing FUS R521H and P525L mutations treated with PARPi, PARPi+ sodium arsenite for 30 minutes, and 30 minutes recovery after PARPi+ Sodium arsenite treatment. Stained with G3BP1 in green, FUS in red and nucleus counter stained with DAPI. Scale bar =20μM for zoomed image scale bar=3 μM, Quantification data represented as mean ± s.e.m derived from three independent experiments, quantification of stress granules was derived from 25 cells. All statistical analysis were performed by two-sided student's-t test using graph pad prism software.

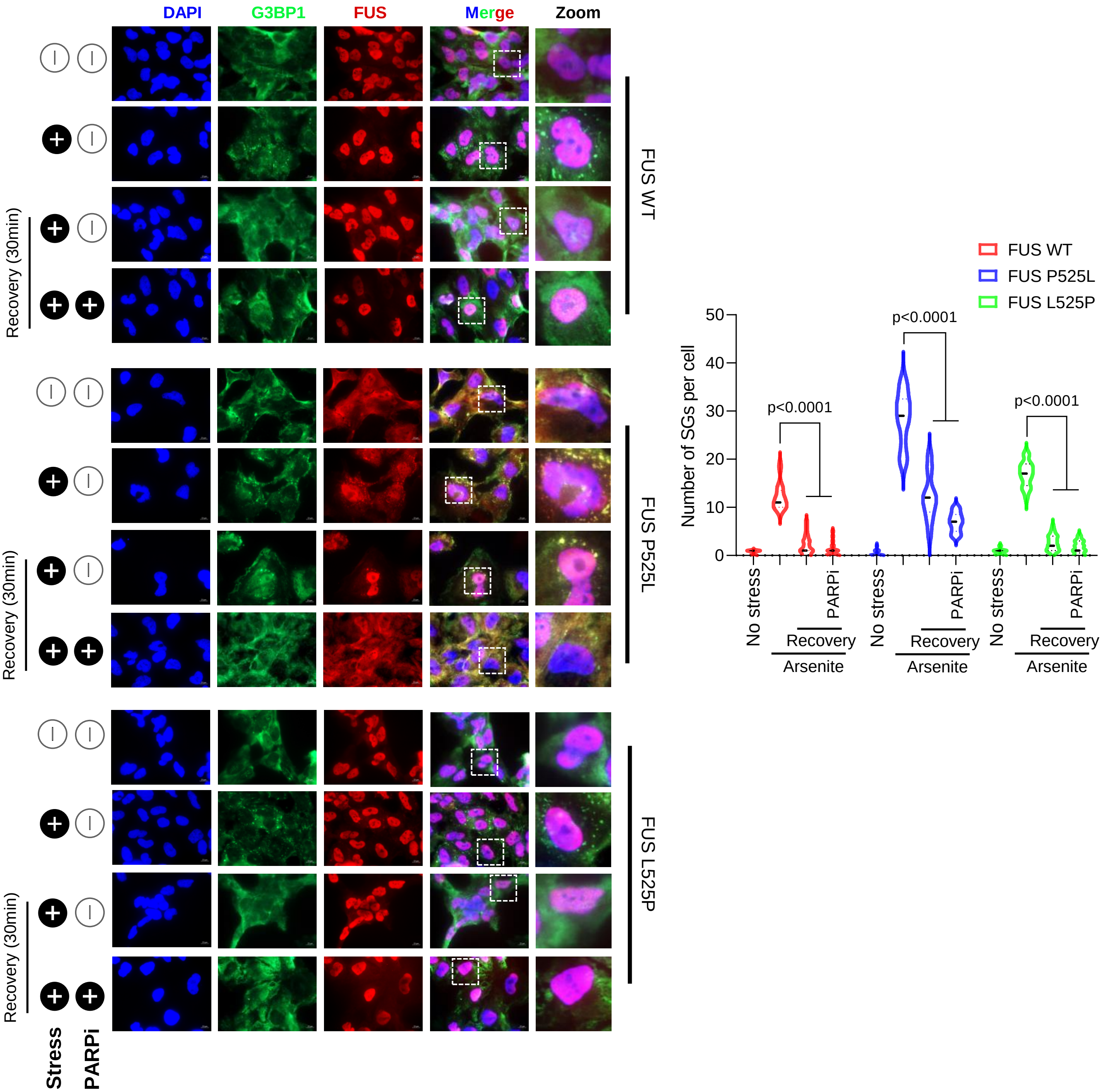


**Supplementary Fig3: FUS pathology associated SGs dynamics alterations can be rescued by correction of FUS mutations**

IF images of control (FUS WT), patient (FUS P525L), and mutation corrected isogenic (FUS L525P) NPSC cells treated with Mock (PBS), Sodium arsenite for 30 minutes, and 30 minutes recovery after Sodium arsenite treatment in presence and absence of PARP inhibitor (Veliparib). Stained with FUS in red, G3BP1 in green and nucleus counter stained with DAPI. Scale bar =10μM, for zoomed image scale bar =3μM. Quantification data represented as mean ± s.e.m derived from three independent experiments, quantification of SGs was derived from 25 cells. All statistical analysis were performed by two-sided student's-t test using graph pad prism software.

**Supplementary Table 1. Primers used in this study**

| | | |
|---|---|---|
| 1 | Human *hPRT* long amplicon Forward primer | 5′-TGGGATTACACGTGTGAACCAACC-3′ |
| 2 | Human *hPRT* long amplicon Reverse primer | 5′-GCTCTACCCTGTCCTCTACCGTCC-3’ |
| 3 | Human *hPRT* short amplicon Forward primer | 5’-TGCTCGAGATGTGATGAAGG-3′ |
| 4 | Human *hPRT* short amplicon Reverse primer | 5′-CTGCATTGTTTTGCCAGTGT-3′ |
| 5 | PARG111 forward primer | 5’-ggccGAATTCATGAATGCGGGCCCCGGCTG-3’ |
| 6 | PARG99 reverse primer | 5’-ggccGAATTCATGATGAGTTCTGTACAAAAAG-3’ |
| 7 | PARG reverse primer | 5’-ggccGAATTCTCAGGTCCCTGTCCTTTGCC-3’ |
